# Computational Evidence for Hierarchically-Structured Reinforcement Learning in Humans

**DOI:** 10.1101/731752

**Authors:** Maria K Eckstein, Anne GE Collins

## Abstract

Humans have the fascinating ability to achieve goals in a complex and constantly changing world, still surpassing modern machine learning algorithms in terms of flexibility and learning speed. It is generally accepted that a crucial factor for this ability is the use of abstract, hierarchical representations, which employ structure in the environment to guide learning and decision making. Nevertheless, how we create and use these hierarchical representations is poorly understood. This study presents evidence that human behavior can be characterized as hierarchical reinforcement learning (RL). We designed an experiment to test specific predictions of hierarchical RL using a series of subtasks in the realm of context-based learning, and observed several behavioral markers of hierarchical RL, such as asymmetric switch costs between changes in higher-level versus lower-level features, faster learning in higher-valued compared to lower-valued contexts, and preference for higher-valued compared to lower-valued contexts. We replicated these results across three independent samples. We simulated three models: a classic RL, a hierarchical RL, and a hierarchical Bayesian model, and compared their behavior to human results. While the flat RL model captured some aspects of participants’ sensitivity to outcome values, and the hierarchical Bayesian model some markers of transfer, only hierarchical RL accounted for all patterns observed in human behavior. This work shows that hierarchical RL, a biologically-inspired and computationally simple algorithm, can capture human behavior in complex, hierarchical environments, and opens the avenue for future research in this field.

## Introduction

Research in the cognitive sciences has long highlighted the importance of hierarchical representations for intelligent behavior, in domains including perception (1), learning and decision making (2, 3), planning and problem solving (4), cognitive control (5), and creativity (6), among many others (7, 8). The common thread across all these domains is the insight that hierarchical representations—i.e., the simultaneous representation of information at different levels of abstraction—allow humans to behave adaptively and flexibly in complex, high-dimensional, and ever-changing environments. Exhaustive non-hierarchical (*flat*) representations, in contrast, are insufficient to achieve human-like behaviors.

To illustrate, consider the following situation. Mornings in your office, your colleagues are working silently, or quietly discussing work-related topics. After work, they are laughing and chatting loudly at their favorite bar. In this example, a context change induced a drastic change in behavior, despite the same interaction partners (i.e., “stimuli”). Hierarchical theories of cognition capture this behavior by positing that we learn strategies hierarchically, activating different behavioral strategies (or “task-sets”) in different contexts.

Although hierarchical representations can incur additional cognitive cost (9), they provide a range of advantages compared to exhaustive flat representations: Once a task-set has been selected (e.g., office), attention can be focused on a subset of environmental features (e.g., just the interaction partner) (10–13). When new contexts are encountered (e.g., new workplace, new bar), entire task-sets can be reused, allowing for generalization (6, 14, 15). Old skills are not catastrophically forgotten (16). In addition, hierarchical representations deal elegantly with incomplete information, for example when contexts are unobservable (6, 17). All these advantages are evident in the current study.

Although we know that hierarchical representations are essential for flexible behavior, how humans create these representations and how they learn to use them is still poorly understood. Here, we hypothesize that learning and using hierarchical representations can be explained under a hierarchical reinforcement learning (RL) framework, in which simple RL computations are combined to simultaneously operate at different levels of abstraction.

RL theory (18) formalizes how to adjust behavior based on feed-back in order to maximize rewards. Standard RL algorithms estimate how much reward to expect when selecting actions in response to stimuli, and use these “action-value” estimates to select actions. Old action-values are updated in proportion to the “reward prediction error”, the discrepancy between action-values and received reward, to produce increasingly accurate estimates. Such “flat” RL algorithms operate over unstructured, exhaustive representations (suppl. Fig. 3A), converge to optimal behavior, are computationally inexpensive, and have led to recent breakthroughs in artificial intelligence (AI) (18).

Broad evidence suggests that the brain implements computations similar to RL: Dopamine neurons generate reward prediction errors (19, 20), and a wide-spread network of frontal cortical regions (21) and basal ganglia (22, 23) represents action values. Specific brain circuits thereby form “RL loops” (17, 24), in which learning is implemented through the continuous updating of action values (11, 25). In this sense, estimating action-values via RL is an algorithm of special interest to cognition: There is strong evidence that the brain implements a simple mechanism to perform the necessary computations.

Nevertheless, RL algorithms have important shortcomings: They suffer from the curse of dimensionality (an exponential drop in learning speed with increasing problem complexity); they lack flexibility for behavioral change; and they cannot easily generalize or transfer old knowledge to new situations. Hierarchical RL (26) mitigates these shortcomings by nesting RL processes at different levels of temporal (27–29) or state abstraction (12, 30).

Recent research has provided support for a plausible implementation of hierarchical RL in the brain: The neural circuit that implements RL is multiplexed, such that distinct RL loops operate at different levels of abstraction along the rostro-caudal axis (10, 24, 31–37). Consistent with this architecture, recent studies have shown signatures of RL values and reward prediction errors at different levels of abstraction in the human brain (29, 38, 39). However, previous studies did not provide evidence that neural signatures of hierarchical value support learning and generalizing hierarchically structured behavior. Thus, it remains unknown whether hierarchical RL indeed supports hierarchical behavior.

The goal of this study is to fill this gap. We investigate hierarchical RL in a novel paradigm that promotes the creation and reuse of hierarchical structure. We provide a fully-fledged computational model that accounts for behavior across a variety of relevant situations: context-dependent learning, context switches, generalization to new contexts, partially-observable problems, and choices at different levels of abstraction. To our knowledge, this is the first study that tests all predictions of hierarchical RL in a single paradigm. Because hierarchical RL makes specific behavioral predictions in each situation, we are able to test the model *qualitatively* against human behavior (40). We then compare our hierarchical RL model *quantitatively* to the two most relevant competing models, flat RL and hierarchical Bayes. The former employs RL, but without hierarchical structure. The latter assumes that high-level decisions are based on Bayesian inference of task-set reliability, rather than RL using task-set values (14).

In the following, we first introduce our hierarchical RL model and experimental paradigm. We then test whether humans show qualitative behaviors that are predicted by the hierarchical RL model, as well as the two competing models. We first show evidence for hierarchical representations in humans, as predicted by both hierarchical RL and hierarchical Bayes, but not flat RL. We employ multiple independent analyses, including switch cost measures and positive and negative transfer. We then provide evidence for human hierarchical value learning, which is only consistent with the hierarchical RL model. We next provide quantitative support for these qualitative results, and show that model comparison supports the hierarchical RL model over flat RL and hierarchical Bayes. The majority of results replicates across three independent participant samples.

## Results

### Computational Models

Our hierarchical RL model is composed of two hierarchically-structured RL processes. The high-level process manages behavior at the abstract level by acquiring a “policy over policies”: It learns which task-set to choose in each context, using “task-set values” (the estimated expected reward of selecting a taskset in a given context). The low-level process acquires these task-sets: it learns which actions to choose in response to each stimulus by estimating “action values” (the estimated expected reward of selecting an action for a given stimulus, within a specific task-set; Fig. 1A).

**Fig. 1.**
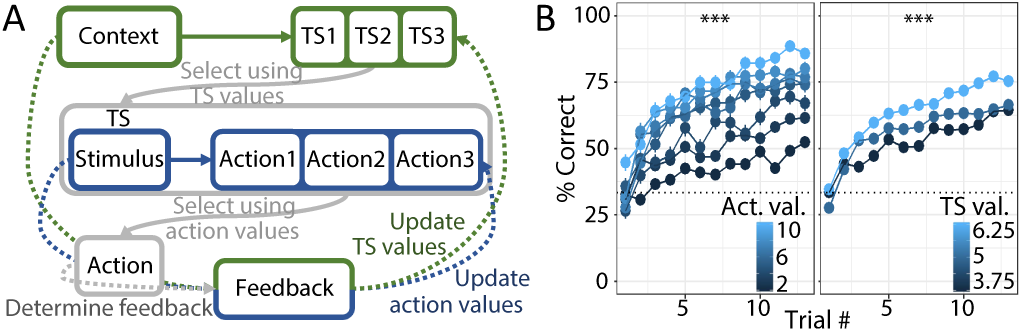
A) Schematic of the hierarchical RL model. A high-level RL loop (green) selects a task-set *TS* in response to the observed context, using TS values. The chosen task-set provides action-values, based on which the low-level RL loop (blue) selects an action in response to the observed stimulus. Task-set and action-values are both learned based on action feedback. B) Human learning curves during the initial learning phase, averaged over blocks. Colors denote underlying action-values (left) and task-set values (right), respectively. Stars show that *both* affect performance (main text), consistent with hierarchical RL. *** indicates *p* < 0.001.

At the beginning of the task, task-sets and actions are picked randomly, but over time, trial-and-error learning leads to the formation of meaningful task-sets, which represent policies that are specialized for particular contexts. Trial-and-error learning also underlies the policy over task-sets that determines which task-set is selected in each context. Thus, our hierarchical RL model is based on two nested processes, which create an interplay between learning stimulus-action associations (low level) and context-task-set associations (high level). Suppl. Fig. 4 shows a step-by-step visualisation of this model.

Formally, to select an action *a* in response to stimulus *s* in context *c*, hierarchical RL goes through a two-step process: (1) It selects a task-set *TS* based task-set values in the current context, *Q*(*TS*|*c*), using 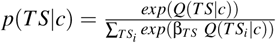. The inverse temperature β_*TS*_ captures task-set choice stochasticity (Fig. 1A). The chosen taskset *TS* provides a set of action-values *Q*(*a*|*s, TS*), which are used to (2) select an action *a*, according to 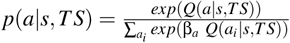, where β_*a*_ captures action choice stochasticity (Fig. 1A; for trial-by-trial behavior, see suppl. Fig. 4B). After executing action *a* on trial *t*, feedback *r*_*t*_ reflects the continuous amount of reward received, which guides learning at both levels of abstraction, i.e., to update the values of the selected task-set and action:

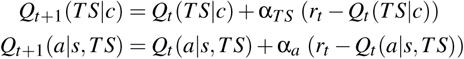

α_*TS*_ and α_*a*_ are learning rates at the levels of task-sets and actions (Fig. 1A; suppl. Fig. 4C).

The flat RL model uses the same mechanism for value learning and action selection, but lacks hierarchical structure: It treats each combination of context and stimulus as a unique state (methods). The hierarchical Bayesian model creates a task-set structure like hierarchical RL, but selects task-sets according to their inferred reliability, rather than task-set values (methods).

### Task Design

We designed a task in which participants learned to select the correct actions for different stimuli (Fig. 2A). The mapping between stimuli and actions varied across three contexts, creating three distinct task-sets (Fig. 2B). Each context appeared in three blocks of 52 trials, for a total of 9 blocks. Contexts differed in average rewards, allowing us to test for RL values at the level of task-sets. After an initial-learning phase of this task (Fig. 2A), participants completed four test phases (Fig. 2C) to hone in on specific predictions of hierarchical RL. Detailed information about the task is provided in Fig. 2, the methods, and supplemental methods.

**Fig. 2.**
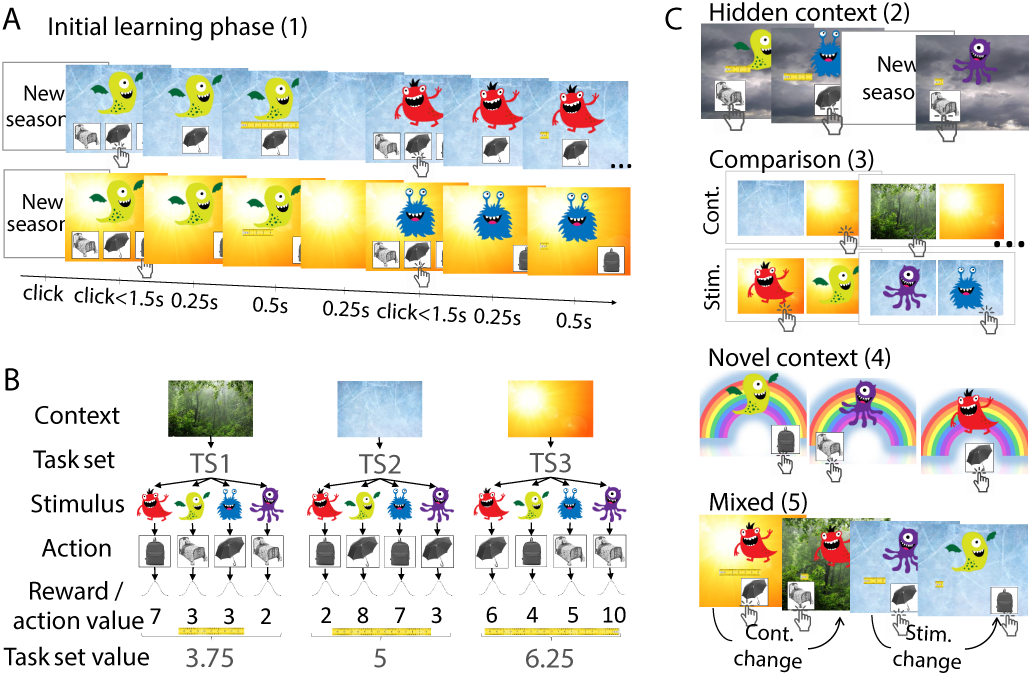
Task design. A) In the initial-learning phase, participants saw one of four stimuli (aliens) in one of three contexts (seasons), and had to find the correct action (item) through trial and error. Each context had a different mapping between stimuli and correct actions, and contexts were presented blockwise. Feedback indicated correctness deterministically, but different context-stimulus-action combinations lead to different rewards (with Gaussian noise). B) Example mapping between stimuli and actions for each context, defining three task-set TS1-3. Average rewards (*task-set values*) differed between contexts. All actions and stimuli had equal average rewards. C) Additional test phases. The hidden-context phase, presented after initial learning, was identical except that contexts were unobservable (season hidden by clouds). This allowed us to test whether participants reactivated previously-learned task-sets. In the comparison phase, participants saw either two contexts (“Cont.”) or two stimuli (“Stim.”) on each trial, and selected their preferred one. We used subjective preferences to assess task-set values (contexts) and action-values (stimuli). The novel-context phase was similar to initial learning, but had a new context and no feedback, to test how participants generalized previous knowledge to new situations. The final mixed phase was similar to initial learning, but not blocked, i.e., both stimuli and contexts could change on every trial, to test for asymmetric switch costs. All test phases were separated by “refresher blocks” similar to initial learning, to alleviate carry-over effects and forgetting.

### Learning Curves and Effects of Reward

As expected, participants’ performance increased within a block, showing adaptation to context changes (Fig. 1B). We also verified that participants were sensitive to continuous differences in reward magnitudes (tape length). RL predicts better performance for larger rewards because these lead to larger action-values, which make correct actions more distinguishable from incorrect ones (see suppl. Fig. 4B for details). Participants indeed showed better performance for high-reward stimuli (Fig. 1B, left).

This effect was predicted by both hierarchical and flat RL. Hierarchical RL additionally predicts better performance for high-valued contexts: Larger rewards create larger reward-prediction errors at the task-set level, which allow for better discrimination between correct and incorrect task-sets, and lead to better task-set selection and performance (see suppl Fig. 4A for details). As predicted, participants also showed an effect of task-set values on performance (Fig. 1B, right).

**Fig. 3.**
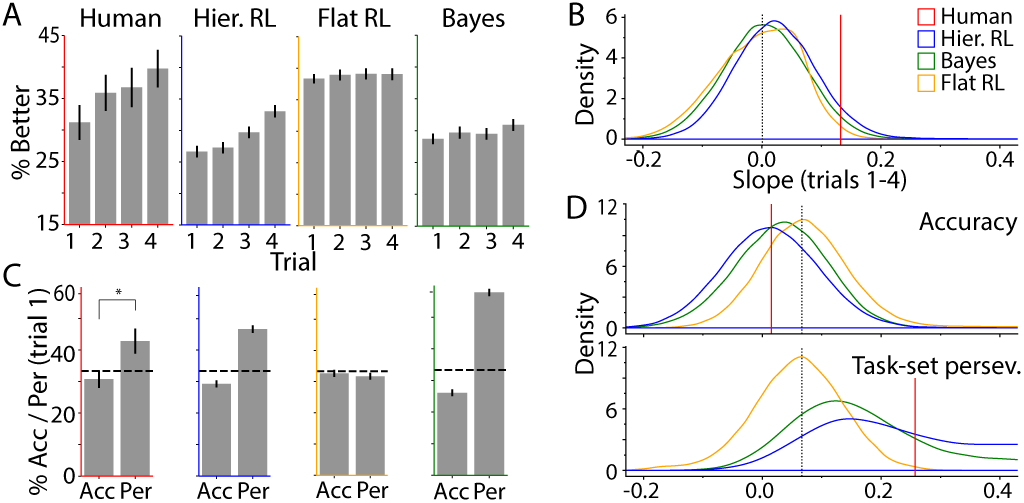
A)-B) Participants reactivated task-sets in the hidden-context phase. A) Human performance (left, red) increased over the first four trials following a context switch, even though different stimuli were presented on each trial. The “best” (methods) simulation based on the hierarchical RL model showed qualitatively similar behavior (blue). The effect was absent in the flat RL model (orange), and present but weaker in the Bayesian hierarchical model (green). B) Slopes of the performance increase in part A), as densities over 50,000 simulations per model, with parameters sampled uniformly at random. These densities approximate marginal model likelihoods for the calculation of Bayes factors. The densities of hierarchical RL and hierarchical Bayes were shifted toward larger slopes, making human-like performance slopes more likely. Dotted line indicates chance. C)-D) Task-set perseveration errors in the initial-learning phase. C) Percent correct trials (“Acc”) and percent task-set perseveration errors (“Per”) on the first trial after a context switch. Humans (left, red): Star denotes significance in repeated-measures t-test. Models: Hierarchical RL and hierarchical Bayes, but not flat RL, qualitatively reproduced human behavior. D) Accuracy and task-set perseveration errors for all simulations, as densities.

**Fig. 4.**
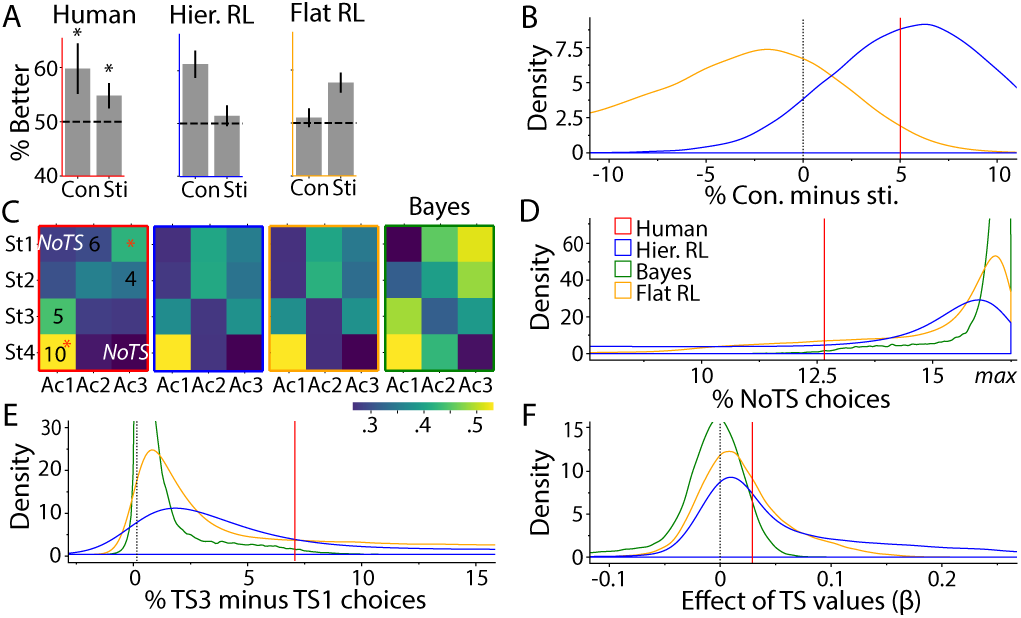
Effects of task-set values on behavior. A)-B) Comparison phase. A) Humans (red) performed better for contexts (“Con”) than stimuli (“Sti”; “% Better”: percentage choosing higher-valued alternative). Stars indicate significant difference from chance (dashed line). The hierarchical RL simulation showed the same qualitative pattern, whereas flat RL showed the opposite. B) Difference between context and stimulus condition, as simulation-based densities. C)-E) Novel-context phase. C) Raw action frequencies. “Ac1-3”: actions; “St1-3”: stimuli. Humans (red frame): Overlaid numbers show actions-values in TS3, the highest-valued task-set, which was chosen frequently. Red stars indicate actions that were correct in multiple task-sets, also selected frequently. “NoTS” indicates actions that were incorrect in all task-sets, selected rarely. Models: Hierarchical (blue frame) and flat RL (orange frame) were qualitatively similar to humans, hierarchical Bayes (green frame) made different predictions. D) NoTS choices in all simulations. E) Difference between percentage of actions consistent with TS3 and TS1. F) Initial-learning phase. Regression weights predicting performance from task-set values, showing that values at both levels affected performance more in the RL models.

To quantify both effects, we conducted a mixed-effects logistic regression model predicting trialwise accuracy from action-values, taskset values, and their interaction (fixed effects), specifying participants, trial, and block as random effects. We approximated action-values as average stimulus-action rewards, and task-set values as average context-task-set rewards, as shown in Fig. 2B. The model revealed significant effects of both action-values, β = 0.38, *p* < 0.001, and task-set values, β = 0.20, *p* < 0.001, on performance (for complete statistics and results in other samples, see suppl. table 1). This provides initial evidence that human choices were sensitive to RL values at two levels of abstraction—actions and task-sets—, as predicted by hierarchical RL.

### Hierarchical Representation

We tested participants’ abstractions in more detail using three independent analyses: switch costs in the mixed phase of the task, reactivation of task-sets in the hidden-context phase, and task-set selection errors during initial learning.

### Asymmetric Switch Costs

Asymmetric switch costs can be evidence for hierarchical representations because changes across trials are more challenging at higher than lower levels of abstraction (41, 42). For example, switching contexts is more cognitively costly than switching stimuli within a context. To test for such asymmetries in our paradigm, we compared trials on which a different stimulus was presented than on the previous trial (but the same context) to those on which a different context was presented (but the same stimulus), using the mixed phase (Fig. 2C). As expected, participants responded significantly slower after context switches than after stimulus switches, *t*(25) = 3.47, *p* = 0.002. This was not due to participants’ initial surprise about the interleaved presentation of contexts in the mixed phase, as the result held throughout the phase (see supplements). Asymmetric switch costs therefore suggest that participants created hierarchical representations, nesting stimuli within contexts, as predicted by hierarchical RL and hierarchical Bayes.

### Reactivating Task-Sets

Did representing the task hierarchically benefit performance, e.g., did it support positive transfer? In the hidden-context phase of our task, contexts were not observable, such that participants could either relearn old stimulus-action mappings from scratch (no transfer), or reactivate previous task-sets, with the correct mappings already in place (transfer). By enabling reactivation of old task-sets, hierarchy has been shown previously to enable better performance and faster learning (6, 14, 34).

If participants reactivated task-sets, we expect a specific pattern of performance in the hidden-context phase, specifically on the first few trials after a context switch, before any stimulus is repeated: Because every trial provides feedback about the appropriateness of the chosen task-set, task-set selection should become more accurate on each trial, and consequently, accuracy should improve.

If, on the other hand, participants did not use task-sets and instead re-learned stimulus-response associations from scratch, as predicted by flat RL, performance can only increase after a stimulus is repeated. Because no stimuli are repeated until the 5*th* trial in our task, the first four trials provide the perfect testing ground to pitch these two predictions against each other, as illustrated in Fig. 3A: Hierarchical RL simulations show increasing performance, whereas flat RL simulations show no change (simulation details in methods and section “Modeling Behavioral Patterns Jointly”).

Human behavior qualitatively matched the predictions of hierarchical RL: Performance increased steadily over the first four trials after a context switch (Fig. 3A), evident in the significant correlation between item position (1-4) and performance, *r* = 0.19, *p* = 0.048. This shows that participants recalled previously-learned stimulus-action mappings rather than relearning them, a signature of task-set transfer.

We next assessed quantitatively which of our three candidate models captured this behavior best. We compared the models using Bayes Factors (*BF*), which we estimated using a method related to Approximate Bayesian Computation (ABC; see methods and supplemental methods (43)). Our method involved simulating synthetic data from each model and estimating the likelihood of human behavior under the simulated data, as illustrated in Fig. 3B. Hierarchical RL surpassed both flat RL, *BF* = 5.12, and also hierarchical Bayes, *BF* = 1.96, in model comparison (suppl. table 3). This confirms the qualitative result, showing that human performance in the hidden-context phase was better captured by hierarchical than flat models.

### Task-set Perseveration Errors

We showed that hierarchy allowed for positive transfer, enabling participants to reactivate old task-sets. However, hierarchy can also lead to negative transfer: When participants select the wrong task-set, the “correct” action according to that taskset is likely to be incorrect in the current context. We call such errors “task-set selection errors”, and focus on a specific subtype, *task-set perseveration errors*. Here, actions are chosen that would have been correct in the previous context, but are incorrect in the current one.

Contrary to flat RL, hierarchical models predict task-set perseveration (methods and example in suppl. Fig. 4A), reflected in high proportions of task-set perseveration errors and low initial accuracy (Fig. 3C and D).

We tested this prediction on the first trial after each context switch during initial learning, and found that participants were more likely to make task-set perseveration errors than to select correct actions, *t*(25) = 2.1, *p* = 0.046, in accordance with hierarchical model simulations (Fig. 3D). Task-set perseveration persisted several trials into the new block, as evident in a logistic regression predicting task-set perseveration errors from trial index (β = −6.83%, *z* = −9.31, *p* < 0.001), task-set values (β = −2.43%, *z* = −1.00, *p* < 0.001), and action-values (β = −14.03%, *z* = −8.45, *p* < 0.001), controlling for block, and specifying random effects of participants.

In summary, the presence of task-set perseveration errors in humans is qualitative evidence for hierarchical processing. Quantitative model comparison supports this conclusion, showing that hierarchical models fit human error patterns better than flat RL (hierarchical vs flat RL: *BF* = 14.99; hierarchical Bayes vs flat RL: *BF* = 10.32; hierarchical RL vs Bayes *BF* = 1.40).

### RL Values at Different Levels of Abstraction

Our results so far focused on hierarchical representations in general, showing that participants created, reactivated, and transferred task-sets. We now test predictions that are unique to hierarchical RL, assessing whether participants acquired RL values at the level of task-sets as well as actions.

### Task-Set Values Affect Subjective Preference

A classic approach to assess RL values in humans is to investigate subjective preferences (44). To investigate whether participants acquired values at both levels, we thus used a comparison phase, where participants selected their preferred out of two items on each trial. Items were either two contexts or two stimuli—testing task-set and action values, respectively (Fig. 2C).

The hierarchical RL model selected contexts based on the task-set values acquired during initial learning, and showed a strong preference for high-valued over low-valued contexts (suppl. Fig. 7A). The flat RL model selected contexts based on average action-values in this context, and showed a much weaker preference (suppl. Fig. 7A). The hierarchical Bayesian model did not track values over contexts and was thus not simulated in this phase.

As predicted by hierarchical RL, participants preferred high-valued over low-valued contexts, *t*(25) = 2.56, *p* = 0.017, indicating RL values at the level of contexts. Quantitative model comparison (Fig. 4B) strongly favored hierarchical over flat RL, *BF* = 1171.65. For completeness, we also confirmed participants’ RL values at the level of stimuli, as predicted by both flat and hierarchical RL, and evident in the preference for high-valued over low-valued stimuli, *t*(25) = 2.11, *p* = 0.045. In conclusion, participants’ preferences were best accounted for by the hierarchical RL model.

We next investigated a different model prediction in the comparison phase: The hierarchical RL model takes two steps to retrieve action-values, but only one to retrieve task-set values. This suggests stimulus selection should be slower and noisier than context selection. Flat RL, on the contrary, takes one step to retrieve action-values, but multiple steps to calculate context-values, suggesting the inverse pattern. Humans showed the patterns predicted by hierarchical RL: RTs were numerically slower and performance was significantly worse for contexts than for stimuli (mixed-effects regression, RTs: β = 148.21, *t*(25) = 1.63, *p* = 0.12, Acc.: β = 0.28, *z* = 2.0, *p* = 0.048; Fig. 4B). Though the effect on RTs did not reach significance here, it was strongly significant in the replication (see supplementary table 1). Quantitative model comparison strongly favored hierarchical over flat RL in terms of accuracy, *BF* = 39.64.

### Task-set Values Affect Performance

As explained above, human initial learning was affected by both action-values and task-set values (Fig. 1B), in accordance with hierarchical RL. To compare our models in this regard, we calculated the effects of task-set values on performance, using a simplified regression model (see suppl. methods). Supporting our qualitative findings, the hierarchical RL model provided a better fit than value-less hierarchical Bayes, *BF* = 6.62, and crucially, than flat RL, *BF* = 1.49 (Fig. 4F).

### Task-set Values Affect Generalization

We showed above that participants preferred high-valued over low-valued contexts (suppl. Fig. 7A). We now test whether participants showed similar task-set preferences in the novel-context phase, that is, when generalizing old knowledge to a new context. For simulations, our hierarchical RL model applied its highest-valued task-set throughout the novel-context phase. The hierarchical Bayes model applied its most reliable task-set. The flat RL model chose actions based on average values (methods).

We labeled each action in the novel-context phase as one of the following: correct in task-set TS3, TS2, TS1, both TS3 and TS1, both TS2 and TS1, or not correct in any task-set (NoTS). Despite the lack of feedback, human participants showed consistent preferences for certain stimulus-action combinations over others (Fig. 4C; see suppl. Fig. 2 for heatmaps of task-set values). They chose NoTS actions less often than other actions, controlling for the frequency of each category, *t*(25) = 2.24, *p* = 0.034. Mappings shared between multiple task-sets (TS2 and TS1; TS3 and TS1) were more frequent than mappings that only occurred in one task-set (TS1, TS2, TS3), controlling for chance level, *t*(25) = 2.83, *p* = 0.0091. This confirms that participants reused old task-sets for new contexts, in accordance with our findings in the hidden-context phase, and prior literature (17). Quantitative model comparison confirmed that the number of NoTS choices was captured better by hierarchical RL than by flat RL, *BF* = 1.78, or hierarchical Bayes, *BF* = 45.60.

Highlighting the role of task-set values, hierarchical RL predicted more actions from the highest-valued TS3 than from the lowest-valued TS1, and a greater difference between the two than flat RL or hierarchical Bayes (Fig. 4E). Humans showed the same pattern, selecting more TS3 than TS1 actions, *t*(25) = 2.58, *p* = 0.016. Bayes Factors confirmed that this difference was captured better by hierarchical RL than flat RL, *BF* = 1.59, or hierarchical Bayes, *BF* = 32.01. Taken together, our hierarchical RL model captured both the reuse of old task-sets in new contexts, and the preference for high-valued over low-valued task-sets.

### Modeling Behavioral Patterns Jointly

Human behavior followed predictions of hierarchical RL qualitatively, and Bayes Factors confirmed quantitatively that this model fit better than the competing ones. However, we treated each behavioral measure independently. We next sought to confirm that it was possible to obtain all behavioral results simultaneously based on a single set of parameters. To this end, we chose one “best” set of parameters for each model (methods), and showed the behavior of this simulation side-by-side with humans, for each behavioral measure. As expected, neither flat RL nor hierarchical Bayes, replicated all qualitative patterns in Figs. 3A, 3C, 4A, and 4C. But importantly, a single set of parameters could capture all qualitative patterns in the hierarchical model. Note that because parameters were not obtained through model fitting, behavior can deviate quantitatively from human data.

## Discussion

The goal of the current study was to assess whether human flexible behavior could be explained by hierarchical reinforcement learning (RL), i.e., the concurrent use of RL at different levels of abstraction (3, 45). We proposed a hierarchical RL model that acquires low-level strategies—or “task-sets”—using RL, and also learns to choose between these task-sets based on RL. We contrasted this model with a flat RL model to highlight the unique contribution of hierarchy, and to a hierarchical Bayesian model, highlighting the contribution of representing learned values hierarchically.

Our hierarchical RL model predicted unique patterns of behavior in a variety of situations. To assess whether humans employed hierarchical RL, we designed a context-based learning task in which multiple subtasks tested these predictions. Indeed, participants’ behavior followed the predictions in all subtasks. The first prediction was that participants would create hierarchical representations. Several independent results supported this claim, including asymmetric switch costs, task-set perseveration errors, and task-set reactivation. These results could not be accounted for by the flat RL model, but were compatible with the hierarchical Bayesian model as well as the hierarchical RL model.

To address the unique aspects of hierarchical RL, we next sought evidence of hierarchical values. Hierarchical RL makes specific predictions about (1) context preferences, (2) effects of contexts on performance, and (3) behavior in new contexts, i.e., generalization. Human behavior showed the predicted patterns in each case: (1) When asked to pick their preferred contexts, participants selected higher-valued ones more often. This suggests that they had formed RL values at an abstract level, in addition to low-level action-values. Participants also performed better when choosing between contexts than stimuli, in accordance with the “blessing of abstraction” of hierarchical representations (46, 47). (2) Task-set values affected performance, with better learning in higher-valued task-sets. This shows that hierarchical representation can explain performance differences between contexts, based on task-set values. (3) When faced with a new context, participants reused previous task-sets, preferring higher-valued over lower-valued ones. This suggests that task-set values guided generalization of old knowledge to new situations.

In summary, human behavior showed all the qualitative patterns predicted by hierarchical RL, but not hierarchical Bayes or flat RL. To quantify the differences between these models, we conducted formal model comparison. Established approaches for model fitting and comparison such as maximum likelihood and sampling-based hierarchical Bayesian methods (48–50) were not applicable in our case, because the model likelihood was intractable. Approximate Bayesian Computation (ABC) and other likelihood-free methods (43, 51, 52) were also infeasible in our setting. We therefore compared models using Bayes Factors, the gold standard of model comparison in Bayesian statics, approximating marginal model likelihoods using simulations, similar to (53). Bayes Factors instantiate an implicit Occam’s razor that accounts for differences in model complexity, such as the larger number of parameters in the hierarchical models compared to flat RL, differences in the functional form of each model, and differences in parameter spaces (53, 54). In this way, Bayes Factors implement a more comprehensive tradeoff between parsimony and goodness-of-fit than traditional methods.

In our paradigm, Bayes Factors showed that hierarchical RL and hierarchical Bayes captured behavioral aspects of *hierarchy* better than flat RL (e.g., task-set reactivation, task-set perseveration), whereas flat RL and hierarchical RL captured *value-based* aspects better (e.g., value-based generalization, effects of values on performance). Furthermore, hierarchical RL uniquely captured the influence of two sets of values on behavior. Overall, Bayes Factors favored the hierarchical RL model over flat RL and hierarchical Bayes. Based on this quantitative confirmation, we next asked whether all results could be jointly observed when simulating the hierarchical RL model with a single set of parameters, to confirm that different parameters were not responsible for different behaviors. We used simulation summary statistics to identify a “best” set of parameters for each model. Only the hierarchical RL simulation qualitatively replicated all human behaviors, but not flat RL or hierarchical Bayes. This shows that seemingly different behaviors, including trial-and-error learning (initial-learning phase), “inference” of missing information (hidden-context phase), subjective preferences (comparison phase), and generalization (novel-context phase), can all be explained in the same overarching hierarchical RL framework.

Note that we have not explored the full space of possible models. In particular, it would be possible to construct a hierarchical Bayesian model that tracks task-set and action-values rather than their reliability, but uses Bayesian inference rather than RL to perform updates. This model might capture the behavioral patterns we observed here. Indeed, our results show evidence for humans’ ability to track values at multiple levels of hierarchy in support of generalizable behavior, but do not speak directly to the exact update process. However, we favor the hierarchical RL formulation of such updates because it is inspired by a rich literature on brain circuits that makes its implementation plausible, and because it is algorithmically simple, with the ability to account for complex cognitive processes.

Many computational models have addressed cognitive hierarchy. How are they related to our model? One important class of hierarchical models is purely Bayesian (7, 55, 56). These models aim to explain, on a computational level of analysis (57), the fundamental purpose of hierarchy for cognitive agents. Our model, on the other hand, is algorithmic, like many pure-RL models: It aims to describe dynamically which cognitive steps humans take when they make decisions in complex environments. Our model is also inspired by the structure of human neural learning circuits (24, 32, 35), thereby extending to the implementational level of analysis.

Some models of hierarchical cognition are method hybrids: Some combine Bayesian inference at the abstract level with RL at the lower level (6, 10). Other, resource-rational models, combine Bayesian principles of rationality with cognitive constrains (58). Frank and Badre (10, 31) proposed a hybrid model that uses Bayesian inference to arbitrate between multiple types of hierarchy and flat RL. In general, hybrid models assume a role for Bayesian inference at higher levels of hierarchy, contrary to our hierarchical RL model. This is an important difference: Hierarchical RL mimics a form of inference (for example, identifying the latent task-set at the beginning of a block; supplementary results 2.1), but cannot do it optimally. It is an important direction for future research to identify whether human behavior is suboptimal in the same way.

Computational models at different levels of analysis (57) are not mutually exclusive. Bayesian inference offers a perspective based on optimality, but it is often intractable and approximations are computationally expensive. RL, on the other hand, uses values to approximate expectations instead of calculating them exactly. Because of its relative computational simplicity, and because it is biologically well supported, RL has often been used as an algorithmic and implementational model. Recent research showed that a neural network implementing hierarchical RL approximated the results of Bayesian inference (17). In other words, hierarchical RL might allow for optimal behavior using simpler computations.

Hierarchical RL was initially proposed in AI (26, 59). A number of AI algorithms has recently been used to model human cognition as well (28, 29, 38, 60, 61), showcasing how intertwined the two fields have become (18, 62, 63). Nevertheless, most hierarchical RL algorithms in AI focus on hierarchy over the time scale of choices (*temporal abstraction*, e.g., breaking up long-term goals into sub-goals). Our hierarchical model, in contrast, focuses on *choice abstraction* (i.e., allowing choice at the level of task-sets and motor actions), a still rare approach in AI (64).

To conclude, classic RL has been a powerful model for simple decision making in animals and humans, but it cannot explain hallmarks of intelligence like flexible behavioral change, continual learning, generalization, and inference of missing information. Recent advances in AI have proposed hierarchical RL as a solution to a number of such shortcomings, and we found that human behavior showed many signs of hierarchical RL, which were captured better by our hierarchical RL model than competing ones.

There is no debate that achieving goals and receiving punishment are some of the most fundamental motivators that shape our learning and decision making. Nevertheless, almost all decisions humans face pose more complex problems than what can be achieved by flat RL. Structured hierarchical representations have long been proposed as a solution to this problem, and our hierarchical RL model uses only simple RL computations, known to be implemented in our brains, to solve complex problems that have traditionally been tackled with intractable Bayesian inference. This research aims to model complex behaviors using neurally plausible algorithms, and provides a step toward modeling human-level, everyday-life intelligence.

## Methods

### Participants

We tested three independent groups of participants, with approval from UC Berkeley’s institutional review board. All were university students, gave written informed consent, and received course credit for participation.

The pilot sample had 51 participants (26 women; mean±age sd: 22.1 ± 1.5), 3 of whom were excluded due to past or present psychological or neurological disorders. Due to a technical error, data were not recorded in the comparison phase for this sample. The second and main sample had 31 participants (22 women; mean age±sd: 20.9±2.1), 4 of whom were excluded due to disorders, and one of whom was excluded because average performance in the initial-learning phase was below 35% (chance is 33%). We added the mixed testing phase for this sample. The third sample had 32 participants (15 women; mean age±sd = 20.8±5.0), 2 of whom were excluded due to disorders. Five participants did not complete the experiment and were excluded when data was missing. The task was minimally adapted for EEG data collection. All statistical tests were conducted in all samples (suppl. table 1 and suppl. Fig. 1), and the supplements discuss sample differences in detail.

### Task Design

Participants first received instructions and underwent the initial-learning phase of the task. The purpose of initial learning was for participants to acquire distinct task-sets, i.e., specific stimulus-action mappings for each context. We also used the initial-learning phase to test for the effects of action-values and task-set values on performance, and to assess errors types predicted by hierarchical RL.

In the beginning, participants were instructed to “feed aliens to help them grow as much as possible”. A tutorial with instructed trials followed, then participants practiced a **simplified task** without contexts: On each trial, participants saw one of four stimuli and selected one of three actions by pressing J, K, or L on the keyboard (Fig. 2A). Feedback was given in form of a measuring tape whose length indicated the amount of reward. Correct actions produced consistent long (mean=5.0) and incorrect actions short tapes (mean=1.0, Fig. 2). When no action was selected, participants were reminded to respond faster next time, and the trial was counted as missed. Participants received 10 training trials per stimulus (40 total), with a maximum response time of 3,000 msec. Order was pseudo-randomized such that each stimulus appeared once in four trials, and the same stimulus never appeared twice in a row.

The **initial-learning phase** had the same structure as training, but stimuli were presented in one of three contexts, each with a unique mapping between stimuli and actions (Fig. 2B). The context remained the same for a block of 52 trials. At the end of a block, a context change was explicitly signaled, before the next block began with a new context. Participants went through 9 blocks (3 per context) for a total of 468 trials. Participants needed to respond within 1.5s, then received reward. Rewards varied between 2-10 for correct actions (Fig. 2B); rewards for incorrect actions remained 1. We chose these numbers to maximize differences between contexts, while controlling for differences between stimuli and actions.

The **hidden-context phase** was identical to initial learning and participants knew they would encounter the same contexts as before, but this time, they were “hidden” (Fig. 2C). There were 9 blocks with 10 trials per stimulus per block (360 total). Context switches were signaled.

The purpose of the **comparison phase** was to assess participants’ subjective preferences for contexts and stimuli, as estimates of their task-set and action-values. Participants were shown two contexts (context condition), or two stimuli in the same context (stimulus condition), and selected their preferred one (Fig. 2C). Participants saw each of three pairs of contexts 5 times, and each of 18 pairs of stimuli 3 times, for a total of 15 + 198 = 213 trials. Participants had 3 sec to respond.

The purpose of the **novel-context phase** was to probe generalization, specifically the reuse of old task-sets in a new context. This phase was identical to the initial-learning phase, except that it introduced a new context in extinction, i.e., without feedback (Fig. 2C). Participants received 3 trials per stimulus (12 total).

The purpose of the final **mixed phase** was to probe switch costs, assessing whether switching contexts was more costly than switching stimuli, indicating hierarchical representation. The mixed phase was identical to the initial-learning phase, except that contexts as well as stimuli could change on every trial. Participants received 3 blocks of 84 trials (252 total), each with 7 repetitions per stimulus-context combination.

To alleviate carry-over effects and forgetting between test phases, we interleaved them with **refresher blocks**, shorter 120-trial versions of the initial-learning phase. More details on task design are provided in the supplemental material.

### Computational Models

We will address in turn how each model behaves in each phase. During **initial learning**, the flat RL model implemented classic model-free (*delta-rule*) RL (18): It treated every combination of a context and a stimulus as a unique state, and learned one RL value for each state and action, as visualized in suppl. Fig. 3A. Using main text notations, values were updated based on *Q*_*t*+1_(*a*|*s, c*) = *Q*_*t*_ (*a*|*s, c*) + α (*r* − *Q*_*t*_ (*a*|*s, c*)), and actions were selected based on 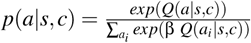.

The flat RL model acquired 36 action-values, based on three parameters α, β, and *f*, whereas the hierarchical RL model acquired 9 task-set-values and 36 action-values (45 total), with six free parameters α_*a*_, α_*TS*_, β_*a*_, β_*TS*_, *f*_*a*_, and *f*_*TS*_ (equations in main text). Suppl. Fig. 3 visualizes the difference between both models, and suppl. Fig. 4 explains hierarchical RL behavior trial-by-trial. The forgetting parameters *f* ∈ [*f*_*a*_, *f*_*TS*_] captured value decay in both models: *Q*_*t*+1_ = (1 − *f*) *Q*_*t*_ + *f Q*_*init*_.

The hierarchical Bayes model also learned task-sets, but acquired their action-values based on correct-incorrect rather than continuous feedback: *Q*_*t*+1_(*a*|*s, TS*) = *Q*_*t*_ (*a*|*s, TS*) + α (*correct* − *Q*_*t*_ (*a*|*s, TS*)). The main difference to hierarchical RL was the selection of task-sets: The Bayesian model chose task-sets based on estimated reliability rather than task-set values, using Bayes theorem to obtain task-set reliabilities: 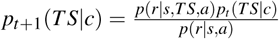, with *p*(*r*|*s, TS, a*) = *Q*(*a*|*s, TS*). Another difference was that hierarchical RL updated *Q*(*TS*|*c*) only for the chosen task-set, whereas hierarchical Bayes kept *p*(*TS*|*c*) up-to-date at all times for all task-sets (6, 14).

Q-values for both models were initialized at the expected reward of chance performance, *Q*_*init*_ = 1.67. The subsequent testing phases started from the Q-values obtained at the end of initial learning.

In the **hidden-context phase**, contexts were not shown, such that models could not directly reuse acquired values that depended on contexts (flat RL: *Q*(*a*|*c, s*); hierarchical RL: *Q*(*TS*|*c*); Bayes: *p*(*TS*|*c*)). All models instead initialized these values at *Q*_*init*_ after each context switch, and then relearned them using the same update equations as before. For flat RL, this resulted in learning an entire new policy *Q*(*a*|*c, s*). For hierarchical models, only high-level information (*Q*(*TS*|*c*) for RL, *p*(*TS*|*c*) for Bayes) had to be relearned, but not action values *Q*(*a*|*s, TS*). This ability to transfer learned values is one of the main advantages of hierarchy.

For the **comparison phase**, we only simulated RL models because the Bayesian model does not provide values at the level of contexts.

To select between two stimuli, RL models first computed the “state value” (18) of each, based on action-values: *V*(*c, s*) = *max*_*a*_ *Q*(*a*|*c, s*) (flat RL) and *V*(*c, s*) = max_*a*_ *Q*(*a*|*s, TS*) *p*(*TS*|*c*), where *p*(*TS*|*c*) = *so f tmax*(*Q*(*TS*|*c*)) (hierarchical RL). Models then selected one stimulus based on a softmax over the two state values. To select between contexts, the hierarchical model repeated the same computation for task-set values: *V*(*c*) = *max*_*TS*_ *Q*(*TS|c*). The flat model, lacking task-set values, used averages over action-values to estimate context preferences on-the-fly: *V*(*c*) = *mean*_*s*_ *V*(*c, s*).

In the **novel-context phase**, models were faced with a context for which they had not learned values. Flat RL used averages over previous action-values to choose: *Q*(*a*|*c*_*new*_, *s*) = *mean*_*c*_ *Q*(*a*|*c, s*). Hierarchical RL [Bayes] applied the previously highest-valued [most reliable] task-set: *Q*(*TS*|*c*_*new*_) = *max*_*c*_ *Q*(*TS*|*c*) [*p*(*TS*|*c*_*new*_) = *max*_*c*_ *p*(*TS*|*c*)].

### Model Comparison

The Bayes Factor *BF* quantifies the support for one model *M*_1_ over another model *M*_2_ by assessing the ratio between their marginal likelihoods, 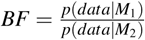. *BF >* 1 provides evidence for *M*_1_. Marginal model likelihoods represent the probability of the data under the model, marginalizing over model parameters θ: *p*(*data*|*M*) =∫ *p*(*data*| *M*, θ) *p*(θ) *d*θ.

For each model, we simulated datasets by drawing model parameters θ uniformly at random. Due to uniform sampling, *p*(θ) is equal for all θ, such that the empirical distribution over simulations approximates the marginal likelihood. To obtain Bayes Factors, we computed the same summary statistics *s*_*m*_ as for humans for each individual simulation (e.g., performance slope in hidden-context phase). We estimated model densities *ŝ*_*m*_ based on a large number of simulations. We obtained marginal model likelihoods as the probability of the human summary statistic *s*_*h*_ under the model, *p*(*s*_*h*_|*ŝ*_*m*_). Bayes Factors are given by 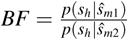.

We drew parameters uniformly at random in a range allowing as broad coverage of possible behavior as possible: 0 < α_*a*_, α_*TS*_, *f*_*a*_, *f*_*TS*_ < 1 and 1 < β_*a*_, β_*TS*_ < 20. Each synthetic dataset consisted of 26 agents simulated on the exact same inputs received by the 26 participants, such that the noise in the synthetic statistics was identical to the one in the human dataset. We simulated 50,000 datasets for each model to assure convergence of the density estimates.

We presented one example datasets for each model in the bar graphs of figures 3A, 3C, 4A. These datasets were obtained by first selecting all of the 50,000 model simulations that fell within a certain range of human behavior for *all* summary statistics (50%-150% for flat and hierarchical RL; 10%-190% for hierarchical Bayes). We then simulated one new dataset per model based on the median parameter values of the selected models. The supplementary methods provide a detailed discussion of our model comparison method and selection of the example datasets.

## Data Availability

All data for this study will be made available for researchers only through the NIMH NDA data base. Analysis and modeling code is available on github: https://github.com/MariaEckstein/TaskSets.

## Acknowledgements

We thank Lucy Whitmore and Sarah Master for their contributions. This project was supported by NIH 1R01MH119383-01 to AGEC.

## Notes

#### Summary of Updates

Thorough rewriting of the manuscript, new analyses, extended supplemental material.

