## Supplemental information for "Computational Evidence for Hierarchically-Structured Reinforcement Learning in Humans"

#### This PDF file includes

Supplementary methods
Supplementary analyses
Figures S1 to S10
Tables S1 to S3

### Supplementary Text, Figures, and Tables

March 23, 2020

### 1 Supplementary Methods

#### 1.1 Participants

We tested our paradigm in three independent samples of participants, replicating most of our major findings. The three versions of the task differed in a few ways: The first sample did not receive the mixed test, and an error in data collection led to the loss of the comparison test data. The second sample received the same task as the first, expect for the addition of the mixed phase. The largest changes—though overall still minor—were necessary for the third sample to enable EEG data collection. The changes concerned mostly timing parameters. Inter-trial intervals were drawn uniformly between 500 and 1.000 milliseconds, in 50 millisecond increments (fixed at 250 milliseconds in previous versions). Intervals before feedback presentation were drawn uniformly between 400 and 800 milliseconds. Testing took place under different conditions, notably in an EEG lab with dimmed light and using a different computer and monitor. Lastly, experimental sessions lasted for 2 hours to accommodate for setting up EEG electrodes on participants' scalps. We chose sample 2 to present in the main text because it is the first sample that includes data from all phases.

Learning curves were qualitatively similar across the samples (suppl. Fig. 1). Overall performance differed slightly, albeit non-significantly, and the large majority of statistical tests replicated across samples (suppl. table 1). In other words, the results reported in the main text were mostly robust to small changes in task design.

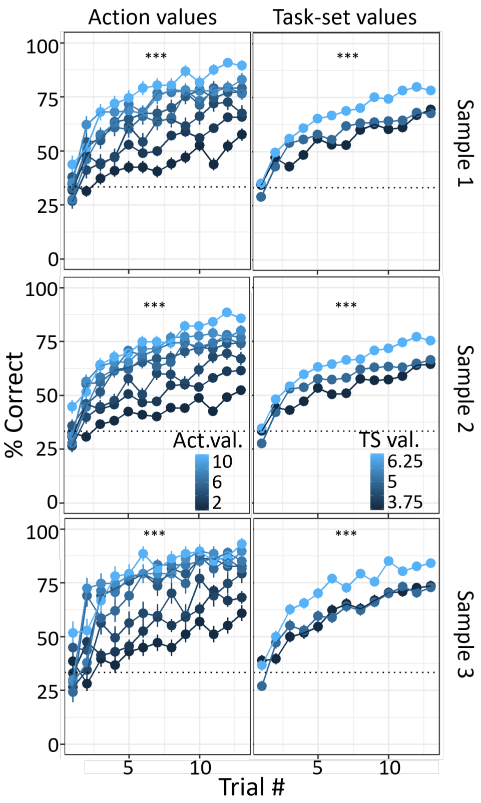

Figure 1: Learning curves of all three samples.

The most notable difference concerned overall task performance of the three samples. EEG participants performed slightly, albeit non-significantly, better, showing qualitatively steeper learning curves (suppl. Fig. 1), numerically better performance and fewer initial selection errors in the initial-learning phase, larger performance increases in the hidden-context phase, and better overall performance in the comparison phase (supple. table 1). It is unclear why performance seemed slightly better in the EEG sample than in the other samples. The most likely reasons include increased attention due to the more involved EEG procedure and changes in timing parameters that slowed the task down slightly.

Table 1 also suggests that task-set values might have had slightly larger effects in the EEG sample than in the other two: In the mixed phase, the effect size of RT switch costs was numerically twice as large, as were the effect size of correlation in the novel-context phase, and differences between stimulus and context condition in the comparison phase, for both accuracy and response times.

Based on these exploratory findings, future research could explore links between task performance and task-set structure.

One test in specific differed between samples: ”More initial selection errors than accurate trials” was statistically significant in the main (second) sample, but not in the first and third (EEG). The most likely explanation were overall performance differences. It is only possible to conduct more mistakes than correct actions when many mistakes are committed. We do not think that this difference in outcomes between samples invalidates any of our claims about hierarchical processing and RL.

Table 1: Statistical tests in the three samples.

| | **Sample 1** No comparison / mixed | **Sample 2** Main sample | **Sample 3** Adapted for EEG |
|---|---|---|---|
| Final sample size (excluded, number and reason) | 48 (3 disorder) | 26 (4 dis., 1 chance perform.) | 30 (2 dis., up to 5 stop early) |
| Mean accuracy during initial learning (sd) | 59.8% (9.3%) | 55.8% (9.3%) | 63.2% (11.5%) |
| **Hierarchical Representation** | | | |
| *Initial-learning phase* | | | |
| Effect of action-values on perf. | $\beta = 0.28(std = 0.04), z = 8.02, p < 0.001$ | $\beta = 0.38(std = 0.05), z = 7.65, p < 0.001$ | $\beta = 0.40(std = 0.05), z = 8.34, p < 0.001$ |
| Effect of task-set values on perf. | $\beta = 0.11(std = 0.04), z = 3.05, p = 0.002$ | $\beta = 0.20(std = 0.05), z = 4.00, p < 0.001$ | $\beta = 0.16(std = 0.05), z = 3.31, p < 0.001$ |
| Interaction between both | $\beta = -0.02(std = 0.01), z = -2.80, p = 0.005$ | $\beta = -0.04(std = 0.01), z = -3.70, p < 0.001$ | $\beta = -0.03(std = 0.01), z = -3.49, p < 0.001$ |
| *Mixed phase* | | | |
| Asymmetric RT switch costs | NA | $t(25) = 3.47, p = 0.002, d = 0.30$ | $t(24) = 4.46, p < 0.001, d = 0.65$ |
| *Hidden-context phase* | | | |
| Reactivating task-sets | $r = 0.12, t = 1.72, p = 0.087$ | $r = 0.19, t = 2.0, p = 0.048$ | $r = 0.34, t = 3.92, p < 0.001$ |
| *Initial-learning phase* | | | |
| Task-set perseverance errors > accurate actions | $t(47) = 0.73, p = 0.47, d = 0.19$ | $t(25) = 2.1, p = 0.046, d = 0.71$ | $t(29) = 0.44, p = 0.67, d = 0.15$ |
| **RL values at different levels of abstraction** | | | |
| *Comparison phase* | | | |
| Stimulus accuracy > 0.5 | NA | $t(25) = 2.11, p = 0.045, d = 0.83$ | $t(28) = 2.58, p = 0.009, d = 0.96$ |
| Context accuracy > 0.5 | NA | $t(25) = 2.56, p = 0.017, d = 1.00$ | $t(28) = 5.61, p < 0.001, d = 2.08$ |
| Context > stimulus acc. | NA | $\beta = 0.28$ ($std = 0.14$), $z = 1.98$, $p = 0.048$ | $\beta = 0.60$ ($std = 0.11$), $z = 5.43$, $p < 0.001$ |
| Context < stimulus RT | NA | $\beta = 148.21$ ($se = 91.14$), $t(25) = 1.63$, $p = 0.12$ | $\beta = 384.90$ ($se = 55.90$), $t(27) = 6.89$, $p < 0.001$ |
| *Novel-context phase* | | | |
| TS3 > TS1 | $t(47) = 3.81, p < 0.001, d = 0.96$ | $t(25) = 2.58, p = 0.016, d = 0.87$ | $t(26) = 2.04, p = 0.052, d = 0.71$ |
| TS2 > TS1 | $t(47) = 3.19, p = 0.003, d = 0.37$ | $t(25) = 1.93, p = 0.065, d = 0.59$ | $t(26) = 1.18, p = 0.25, d = 0.38$ |
| TS3and1 > TS2and1 | $t(47) = 3.14, p = 0.003, d = 0.59$ | $t(25) = 1.37, p = 0.18, d = 0.30$ | $t(26) = 3.82, p < 0.001, d = 1.07$ |

#### 1.2 Task Design

##### 1.2.1 Additional Task Information, Including Randomization

The association between task-sets and contexts was randomized between participants; for example, the winter context might be associated with the highest-valued TS3 for participant 1, but with the lowest-valued TS1 for participant 2. Similarly, the association between stimuli and alien characters, and between actions and items was randomized. The position of actions, i.e., the keyboard key associated with it, were not randomized between trails. In other word, the bed, umbrella, and backpack always appeared in the same position on the screen within one participant, but they differed between participants.

We randomized the mapping between contexts (e.g., winter) and their role (e.g., highest-valued task-set) to avoid systematic biases in participant response, e.g., due to the semantics of different objects. We also conducted basic analyses to confirm that specific objects did not lead to systematic response biases.

Trial order was randomized in the following way for the initial-learning phase and hidden-context phase: Each phase consisted of several blocks. Each block included a single context (season), but all stimuli (aliens). The order of context blocks within each phase was pseudo-randomized such that each context block appeared once within a macro-block of three blocks, and the same context never appeared twice in a row. Context changes were signaled explicitly, even though changes in the context were visually very salient. This was done to avoid mistakes based on uncertainty about the current context. Both the initial-learning phase and the hidden-context phase consisted of three macro-blocks (nine blocks total).

Stimulus order within each block was pseudo-randomized in a similar way: A mini-block consisted of the four stimuli randomized in order, and mini-blocks were combined such that the same stimulus never appeared twice in a row. This randomization ensured that each stimulus was presented equally often in each position across a block. Thirteen mini-blocks formed one block for the initial-learning phase, and ten for the hidden-context phase, for a total of 9 (blocks) * 13 (mini-blocks) * 4 (stimuli) = 468 trials in the initial-learning phase, and 9 * 10 * 4 = 360 trials in the hidden-context phase.

The novel-context phase only contained a single block because it introduced a single new context (rainbow). This block consisted of 3 mini-blocks, with stimuli randomized as before, for a total of 3 (mini-blocks) * 4 (stimuli) = 12 trials. No feedback was given. The low number of trials was chosen to limit the risk of participants disengaging in the absence of feedback.

The mixed phase was structured slightly differently. It consisted of mini-blocks of 12 trials (one for each combination of stimuli [4] and contexts [3]). Trial order was randomized within each mini-block. Seven mini-blocks were combined into one block, and self-paced breaks separated three blocks in total, for a total of 3 (blocks) * 7 (mini-blocks) * 12 (items per mini-block) = 252 trials. The correct mappings between contexts, stimuli, and actions were the same in the mixed phase as before during initial learning and in the hidden-context phase, and participants received the same kind of feedback as before. The only difference was that contexts were no longer presented blockwise, and both stimuli and contexts were allowed to switch on every trial.

The comparison phase reused the same objects as before, but presented participants with a different task: Instead of selecting an action for a given stimulus (and context), participants saw two different stimuli (and a context) on the screen and had to select their preferred one, via button press ("stimulus condition"). The context was always the same for two stimuli that were presented together, to facilitate the task for participants as well as choice analysis.

We counted a trial as correct in this test when participants chose the stimulus that had led to larger reward during initial learning (larger action-value). For example, presented with the red and the purple alien in the rainy season, a correct choice would be to pick the red alien because it had led to a reward of 7, and not the purple alien, which had led to a reward of 2 (Fig. 2B).

The context condition was similar to the stimulus condition, except that participants saw just two contexts without any stimuli, and selected their preferred one. We counted an action as correct in this condition when participants selected the context with the larger average reward (task-set value), as shown in 1B). We used the stimulus condition of this phase to test for the formation of action-values in our participants, and the context condition to test for task-set values.

Trial order in the comparison phase was randomized using a block structure like before. Each block in the context condition consisted of all pairs of two contexts, i.e., 2 choose 3 [contexts] = three trials. Each block in the stimulus condition consisted of 2 choose 2 [stimuli] = six trials for each context, i.e., 6 (pairs) * 3 (contexts) = 18 trials in total. Trials were randomized within each block, and blocks were combined such that the same trial did not repeat twice in a row. The context condition had five blocks (5 * 3 = 15 trials total) and the stimulus condition 3 (3 * 18 = 54 trials total).

In addition to the context and stimulus condition just described, we also tested an item condition (presenting each pair of two items together), a "pure" stimulus condition (only stimuli, without contexts), and a "mixed" stimulus condition (two different stimuli, with two different contexts). These conditions were not of interest and presented after the context and stimulus condition.

##### 1.2.2 Task Timing

We limited response times to 1.5 seconds. Multiple considerations went into this decision. First, this is a usual task timing for most reinforcement learning experiments, and keeping the timing similar allows for comparison between them. Another, more pragmatic, reason was to keep the experiment within 60 minutes to limit participant fatigue, while ensuring a sufficient number of trials in each phase. Last, we aimed to motivate participants to employ reinforcement learning rather than cognitive control or effortful strategizing, which require more time.

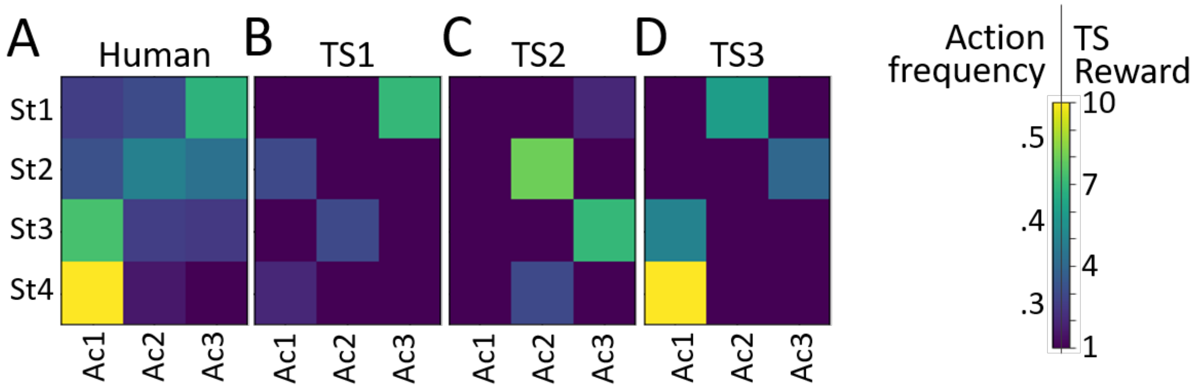

Figure 2: A) Raw human action probabilities in the novel-context phase. B)-D) Action-values of the three task-sets.

###### 1.2.3 Details about Task-Sets

The task-sets were constructed such that there was only one correct action for each stimulus in each context, but the same action could be correct for multiple stimuli. E.g., the bed was the only correct action for the yellow alien in the rainy context in the example shown in 2B. Selecting the bed for the yellow alien in this context led to a measuring tape of length 3. The backpack and the umbrella were both incorrect, and selecting either of these led to a tape of length 1. In the same context, the bed was also the correct response for the purple alien, for which the reward was a tape of length 2.

To obtain action-values and task-set values in Fig. 2B, we assessed the average rewards for a correct response. For example, choosing the backpack for the red alien in the rainy context was rewarded with measuring tapes of length 7±noise, averaging out to an expected reward of 7. Similarly for task-set values, we averaged the rewards of correct actions in each task-set (Fig. 2B). When we indicate ”higher-valued” task-sets, we refer to task-sets that have larger task-set values thus calculated.

The heatmaps in suppl. Fig. 2 show the three task-sets visually. The information is identical to what is presented in 2B. Task-set values are shown side-by-side with human raw action frequencies in the novel-context phase (replicated from Fig. 4C).

##### 1.3 Regression Models in Humans and Simulations

To assess the effects of action-values and task-set values on performance, we calculated regression models predicting performance from both, and assessed their respective regression weights. To analyze human data (Fig. 1B), we ran a mixed-effects regression model that controlled for a range of other factors as well (e.g., block, item position), as explained in the main text. To estimate model likelihoods, we had to get a similar measure from our computational models. Nevertheless, running the mixed-effects model we used for humans in all 150,000 model simulations was infeasible due to the computational demands. We therefore ran a simpler regression model in simulations, predicting performance from just action-values and task-set values. We present the regression weights of task-set values obtained from these models in the model distributions (Fig. 4F). For model comparison only, we re-ran the simplified regression model on human data as well; the results of the simplified model are shown in Fig. 4F (red line), and were used to calculate the model likelihoods for this measure.

##### 1.4 Computational Models

###### 1.4.1 Values in Flat and Hierarchical RL

The hierarchical RL model had nine task-set values, which specify the value of each task-set (3) for each context (3). The number of three task-sets was chosen to accommodate for the three contexts, and is consistent with previous research on how many task-sets humans entertain in parallel (1).

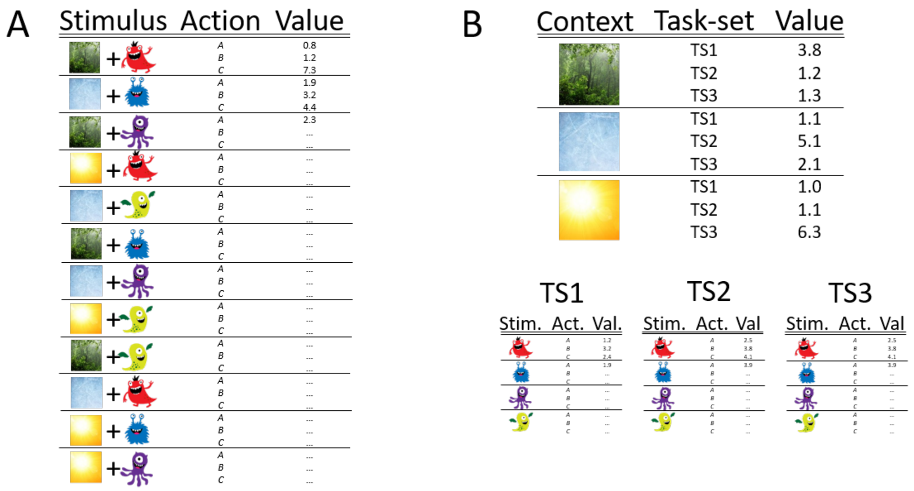

Figure 3: Visualization of learning in flat and hierarchical RL. A) Flat RL learns an ”exhaustive” value table, treating each combination of context and stimulus independently from each other. B) Hierarchical RL learns distinct ”task-sets”, which contain stimulus-action values or ”low-level values” (bottom). Task-sets are associated with contexts using a table of task-set or ”high-level” values (top).

In addition, the hierarchical RL model contains 3 (task-sets) * 4 (stimuli) * 3 (actions) = 36 action-values, specifying the value of each action for each stimulus, in each task-set. The flat RL model only contains 3 (contexts) * 4 (stimuli) * 3 (actions) = 36 values in total.

##### 1.4.2 Visualization of Flat and Hierarchical RL

Flat RL learned independent action-values for each context-stimulus-action combination, which can be visualized in a single ”flat” value table (suppl. Fig. 3, left). Hierarchical RL learned values for each context-task-set combination and for each task-set-stimulus-action combination, therefore requiring two separate value tables (suppl. Fig. 3, right).

##### 1.4.3 Trial-By-Trial Behavior of a Hierarchical RL Agent

To shed light on the processes that underlie trial-by-trial choices, we zoomed in on the behavior of a hierarchical RL agent in the initial-learning phase (suppl. Fig. 4). Agent behavior showed several interesting patterns, for example better performance in high-valued than low-valued contexts, just like humans. Unlike in human participants, we were able to assess the RL process directly in the simulated agent, allowing us to investigate the precise dynamics that gave rise to this pattern.

The most striking behavioral differences between contexts arose for task-set selection: While the highest-valued context was in place (suppl. Fig. 4A, red), the agent selected the same (red) task-set throughout. During the lowest-valued context (green), on the other hand, the agent circled through all three task-sets inconsistently. An intermediate amount of task-set switching occurred in the middle-valued context (blue). Consistent task-set selection therefore went hand-in-hand with better performance. The reason for this was that consistent task-set selection allowed the agent to use feedback maximally efficiently: Consistent task-set selection means that every feedback is used to update the action-values of the same task-set, which therefore quickly turns into an optimal strategy for this context (suppl. Fig. 4B, action-values inside the red box). In contrast, inconsistent task-set selection was related to poor performance because it led to suboptimal use of feedback: Action-value updates were applied to all task-sets, which impeded the creation of a single optimized task-set (green box), and perturbed action-values of other, already-optimized, task-sets.

Differences in task-set switching ultimately arose from differences in reward sizes between contexts (Fig. 2B). Mechanistically, large rewards quickly led to large task-set values and consistent task-set selection. Consistent task-set selection allowed for more efficient use of feedback and the formation of optimized action-values. Optimized action-values, in turn, enabled correct action selection and led to rewards, which further increased task-set values, etc. Small rewards had the opposite effect, leading to small task-set values, frequent task-set switching (suppl. Fig. 4D), suboptimal action-values, lack of rewards, etc.

The performance differences exemplified by this agent are a general feature of hierarchical RL. Hierarchical RL also showed the other behaviors observed in humans, such as task-set perseveration errors (suppl. Fig. 4A), quick task-set reactivation after context switches (in initial learning and hidden-context phase), and effects of reward size on performance (suppl. Fig. 4).

##### 1.4.4 Model Comparison

**Model Comparison is Relative** The goal of model comparison is to test which one of two (or more) models explains a given dataset better. As such, model comparison is always relative, and does not provide an "absolute" measure of model fit. The quality of model comparison depends on the models included in the comparison, and on the supporting analyses that are performed externally to model comparison. We compared our hierarchical RL model to a flat RL and a hierarchical Bayesian model to test both of its unique aspects, aiming to provide as meaningful a model comparison as possible. We also supported our modeling analyses with qualitative behavioral analyses, and show that human behavior exhibited specific patterns that were only predicted by hierarchical RL, but not the other models.

**Model Comparison Method** We explained in the main text that our hierarchical RL model could not be fitted using traditional parameter fitting methods, such that we had to chose an alternative method for model comparison, namely simulation-based Bayes Factors.

We took great care to ensure that our results were not based on our chosen hyper-parameters. Specifically, we explored different ranges of $\beta$ parameters, the only parameters whose range was not given naturally. Changing these ranges had slight effects on the distributions of the models, but did not affect Bayes Factors in a meaningful way, and therefore did not affect our conclusions. We also explored different numbers of simulations per model, and found that this number had no noticeable effects beyond a reasonably high number of a few thousands. We also explored different values of the range parameter when selecting human-like simulations (suppl. Fig. 5A). The results were highly consistent for different ranges.

An overview of the results of model comparison for all measures is provided in table 3.

Table 2: Bayes factors. Numbers greater than 1 support the model mentioned first (usually hierarchical RL), numbers smaller than 1 support the model mentioned second.

| | HRL vs flat | HRL vs Bayes | Bayes vs flat |
|---|---|---|---|
| **Hierarchical representation** | | | |
| Hidden-context phase, slope (Fig. 3B) | 5.12 | 1.96 | 2.61 |
| Initial-learning phase, accuracy $1^{st}$ trial (Fig. 3c) | 2.88 | 1.31 | 2.21 |
| Initial-learning phase, task-set perseverance $1^{st}$ trial (Fig. 3c) | 14.49 | 1.40 | 10.32 |
| **RL values at two levels of abstraction** | | | |
| Comparison phase, stimulus accuracy (suppl. Fig. 7B) | 0.48 | NA | NA |
| Comparison phase, context accuracy (suppl. Fig. 7A) | 1171.65 | NA | NA |
| Comparison phase, cont. minus stim. acc. (Fig. 4B) | 39.64 | NA | NA |
| Initial learning, regr. TS values on perf. (Fig. 4F) | 1.49 | 6.62 | 0.22 |
| Novel-context phase, frequency NoTS choices (Fig. 4D) | 1.78 | 45.60 | 25.55 |
| Novel-context phase, TS3 minus TS1 chocies (Fig. 4E) | 1.59 | 32.01 | 20.14 |

##### 1.4.5 Selection of Example Model Simulations

We presented one example simulation from each model in the bar graphs of figures 3A, 3C, and 4A. We obtained these simulation results in two steps: We first defined performance criteria around human behavior (see below) and selected a small subset of the 50,000 simulations that we had created for each model that matched these criteria. We then calculated the median parameter values across the selected simulations for each model, which are shown in table 3. We used the obtained parameters to create a new simulation for each model. The re-simulation of an independent dataset avoids problems of double-dipping and biased selection that would arise if an already-simulated dataset was presented based on its preferable performance.

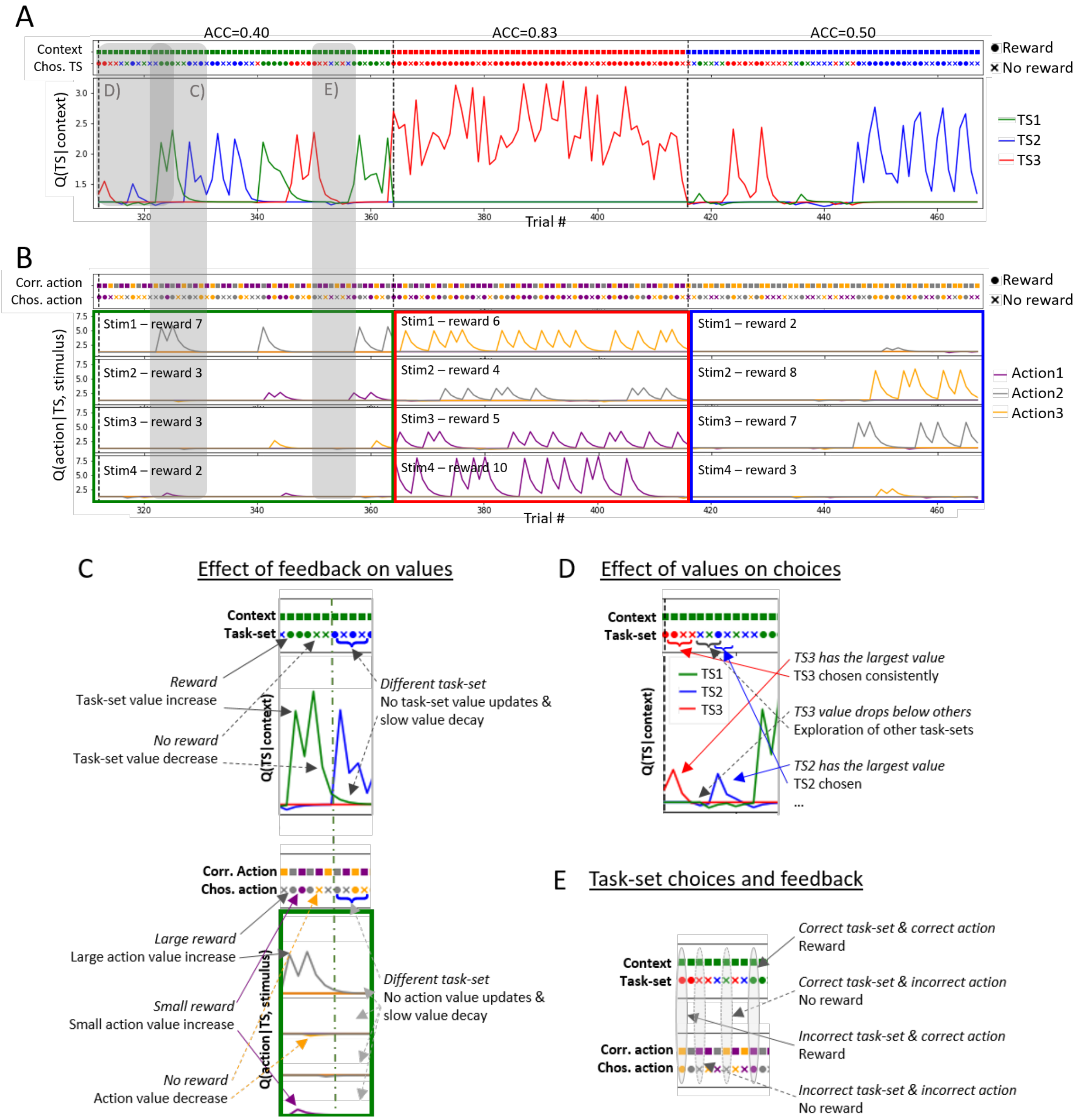

Figure 4: Trial-by-trial behavior a hierarchical RL agent from the population shown in Figures 3 and 4. A) RL at task-set level. Top: Sequence of contexts and agent's selected task-sets. There was no imposed mapping between contexts and task-sets; identical colors highlight which task-sets became specialized for each context during training. Bottom: Evolution of task-set values. B) RL at action level. Top: Sequence of stimuli (represented by correct actions) and agent's selected actions. Bottom: Action-values over time. Only the specialized task-set is shown for each context, indicted by box color. Task-sets contain action-values for all four stimuli (Stim1, Stim2, etc.). C) Top (bottom): Task-set (action) values were affected by reward (increase), no reward (decrease), and non-selection (slow decay, i.e., forgetting). D) Probability of task-set selection was based on relative task-set values. Agents circled through task-sets when no task-set was optimized and rewards were rare. E) Feedback reflects the validity of action selection, not of task-set selection. Task-set values arise from an indirect process mediated through action-values.

Table 3: Median parameters of all models selected for similarity to human behavior. These parameter values were used to simulate a single new dataset to show side-by-side with humans

| | $\alpha_a$ | $\beta_a$ | $f_a$ | $\alpha_{TS}$ | $\beta_{TS}$ | $f_{TS}$ |
|---|---|---|---|---|---|---|
| Hierarchical RL | 0.49 | 9.74 | 0.47 | 0.29 | 13.77 | 0.17 |
| Flat RL | 0.49 | 10.11 | 0.31 | NA | NA | NA |
| Hierarchical Bayes | 0.79 | 13.28 | 0.21 | NA | 12.08 | 0.44 |

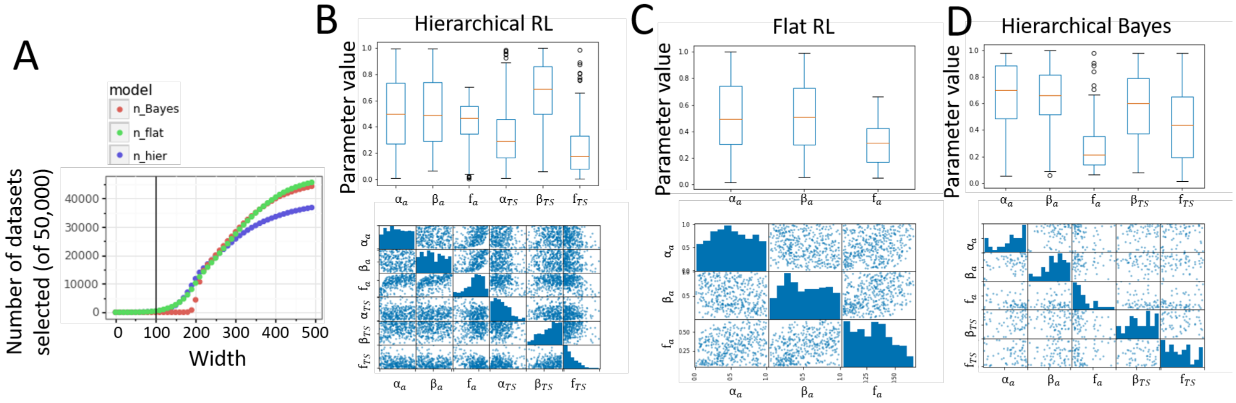

Figure 5: A) Number of selected simulations for each model, dependent on the chosen range around human performance. B)-D) Box plots and correlation matrices for the parameters of the selected datasets. Parameter values of $\beta_a$ and $beta_{TS}$ are scaled down by a factor of 20 for easier comparison. All other parameters naturally lie in the range between 0 and 1. B) Hierarchical RL, C) Flat RL, D) Hierarchical Bayes.

We defined the following criteria to select model subsets. We took note of human behavior for each of our summary measures (e.g., amount of NoTS choices in novel-context phase, accuracy in the context condition of the comparison phase, etc.). We then calculated a range around human performance for each summary measure, and selected all simulated datasets that fell within the ranges *of all measures simultaneously*. Fig. 5A shows how many simulations were selected from each model based on the range around human performance.

We chose a window width of 100 around human measurement to select hierarchical and flat RL simulations, and width 180 to choose hierarchical Bayesian simulations. At width 100 (50%-150% of human performance), 314 / 50,000 (0.63%) hierarchical RL simulations were selected, and 434 / 50,000 (0.87%) flat RL ones (0 hierarchical Bayes datasets). At width 180 (10%-190% of human performance), 65 (0.13%) hierarchical Bayes simulations were selected.

We next inspected the parameters of the selected simulations in each model. Parameters were no longer distributed uniformly (suppl. Fig. 5B-D) like in the source distribution of the initial 50,000 simulations per model. This shows that for each model, human-like behavior was more likely under some parameters than others. For example, human-like performance in the hierarchical RL model was more likely to arise with low forgetting of task-set values - this can be interpreted as relating to participants' ability to reuse task-sets, which would not be the case if task-set values were fully forgotten.

Simulating new datasets in this way avoids problems of double-dipping when presenting example model behavior. But despite its conceptual similarity to ABC rejection sampling (2), this method is not a parameter fitting procedure, i.e., the parameters in table 3 should not be interpreted as "best fits to the human data". One aspect that is lost by this procedure, for example, is the strong interdependence between parameters, for example relationships between $\beta_a$ and $f_a$ in hierarchical RL (suppl. Fig. 5B-D, bottom). The parameters obtained in this way were a basic estimate of reasonable parameter settings at the group level, and were merely meant to highlight the qualitative patterns that can arise from each model.

### 2 Supplementary Analyses

#### 2.1 Additional Analysis of Initial Learning

Most of our analyses in the main paper followed the same logic: We made a behavioral prediction based on the hierarchical RL model, revealed the predicted pattern in human behavior, and concluded that hierarchical RL captured human behavior in qualitative terms. Here, we report an analysis that tests a prediction based on the hierarchical Bayesian model, to provides an inverse test.

Some problems can only be solved by Bayesian inference, but not through RL (in its basic form). For example, a Bayesian model is able to switch to correct behavior immediately after receiving a single diagnostic feedback, whereas RL needs to try out many actions to achieve this (in the right set-up). We therefore tested for markers of immediate switching in human data, as evidence for the hierarchical Bayesian model, and against hierarchical RL.

Nevertheless, we found no evidence for such behavior in our task. To test this, we first selected diagnostic and non-diagnostic trials in the hidden-context phase, and then compared participants' performance in the trials immediately following these trials. We defined diagnostic trials as trials in which feedback (correct vs. incorrect) indicated the correct task-set with certainty, such as receiving a reward after selecting the umbrella for the blue alien, which is only correct in TS1. Non-diagnostic trials were defined as trials in which the was not the case, e.g., receiving a reward after selecting the backpack for the red alien is correct in both TS1 and TS2 (even though reward amounts differ; see below). Most correct trials were diagnostic, whereas all incorrect trials were non-diagnostic in this sense (e.g., not receiving a reward after selecting the umbrella for the green aliens still leaves both TS1 and TS3 as possible candidates). We therefore restricted our analysis to correct trials only.

We found no difference in performance between dignostic and non-diagnostic trials (hidden-context phase, performance on trials immediately following diagnostic trials: 68.0%; performance on trials immediately following non-diagnostic trials: 66.6%; difference between the two in repeated-measures t-test: $t(25)=0.60$, $p=0.56$). This shows that we were unable to find the behavior predicted by the hierarchical Bayesian model in our task, consistent with our hypothesis that human cognition employs hierarchical RL rather than Bayes.

Nevertheless, our task was not designed to test this prediction specifically, and the test just described had one potential confound, so we defer from drawing definite conclusions from it. The potential confound is that all correct feedback in our task indicates the correct task-set with certainty, not just the trials we termed diagnostic above: Even though some stimulus-action mappings are shared between task-sets, their rewards always differ. For example, action 3 is correct in both TS1 and TS2 for alien 1 (Fig. 2B). But because the reward is 7 for TS1 but 2 for TS2, correct feedback indicates with certainty which one is in place.

Thus, an agent with a perfect model of the task could, in theory, know with certainty which TS to select after a single reward. Whether our discrimination of diagnostic and non-diagnostic trials is valid therefore depends on whether humans have such a perfect model of our task. To conclude, our results suggest that participants were not able to quickly switch to a correct task-set after a single diagnostic feedback, despite the fact that the structure of the task could allow such. Instead, their learning process shows a slower trajectory, consistent with our hierarchical RL model rather than perfect inference.

#### 2.2 Additional Analyses of Mixed Phase

##### 2.2.1 Basic Behavior

Average performance in the mixed phase was 50.3% (sd=20.0%), as compared to 55.8% (sd=9.3%) during initial learning (chance=33.3%). The numerically lower performance might reflect the increased difficulty of the task when both stimuli and contexts were allowed to change on every trial. Nevertheless, differences between action-values and task-set values persisted. As expected, learning within blocks was not evident (suppl. Fig. 6A).

##### 2.2.2 Switch Costs

Blocking contexts during initial learning might induce expectations in participants that contexts are necessarily blocked. The slower RTs after context switches than stimulus switches in the mixed phase could be a result of a violation of participants' expectation thus formed, rather than an index of hierarchical structure. We took several measures to alleviate this concern. First, participants were told explicitly at the beginning of the mixed phase that contexts would change quickly and unpredictably:

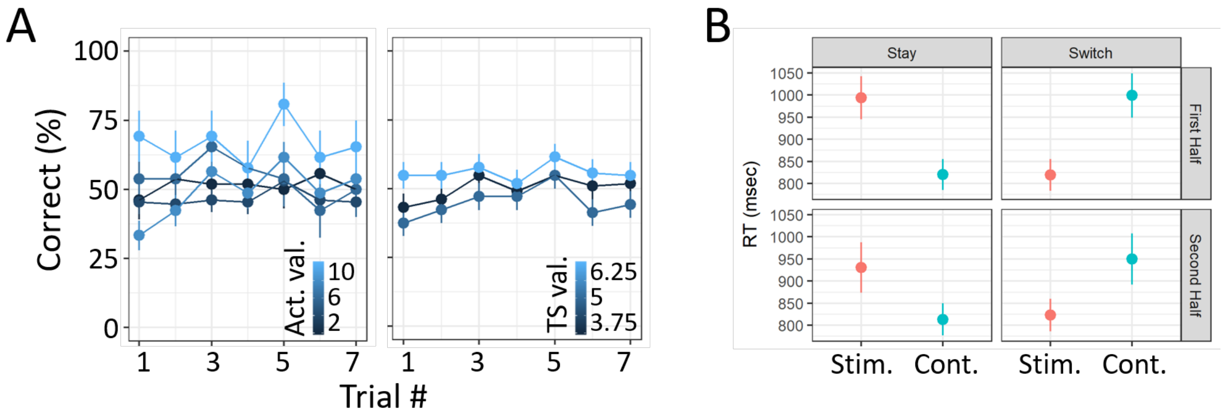

Figure 6: Mixed phase behavior. A) Performance broken up by action-values ("Act. val.", left panel) and task-set values ("TS val.", right panel). B) Response times on correct trials, in the mixed phase. The left panel shows stay trials, i.e., when the same stimulus ("Stim.", red) or context ("Cont.", blue) is repeated. The right panel shows switch trials. The top and bottom panel show the similarity between the first and second half of the mixed phase.

"You will next encounter the *chaotic season*. In the chaotic season, the weather changes very quickly. It can be rainy one day, and then sunny the next."

Nevertheless, expectations might persist implicitly. To investigate whether this was the case, we compared the RT effect in the first and second halves of the mixed phase. Our reasoning was that expectations about trial order should fade away quickly once participants realize that contexts are presented in random order. Therefore, RT effects should diminish over time. If the RT effects were caused by participants' hierarchical representation, on the other hand, the RT effect should persist.

We split the mixed phase into two halves of equal size, and tested for the RT effects in both halves separately. The RT effect was present in both, with no significant difference between them (suppl. Fig. 6B); all trials: t(25)=3.5, p=0.002; only first half: t(25)=2.5, p=0.02; only second half: t(25)=2.7, p=0.01; difference between first and second half: t(25)=0.9, p=0.40). These results suggest that expectations formed by blocked context presentation were unlikely a complete explanation of the RT effects. The more likely explanation was the hierarchical representation of stimuli within contexts.

#### 2.3 Additional Analysis of Comparison Phase

In the main text, we only showed the performance difference between stimulus and context condition, but not raw performance in each individually. Suppl. Fig. 7 and table 2 provide this information. As shown in the figure, the hierarchical RL model was likely to obtain better performance in the context condition (Fig. 3A), but worse performance in the stimulus condition, compared to flat RL. Bayes factors therefore favored hierarchical RL over flat RL in the context condition, $BF = 1.31$, but flat RL over hierarchical RL in the stimulus condition, $BF = 0.79$.

#### 2.4 Relationship Between Model Parameters and Hierarchical Behavior

The behavioral predictions differed qualitatively between our tested models: E.g., the hierarchical RL model predicted that performance would improve during the first four trial of the hidden-context phase, whereas the flat RL stated that performance could not yet improve. Despite this qualitative difference in prediction—the presence versus absence of an effect—, the observable behavioral pattern—i.e., the slope over performance in the first four trials—lay on a continuum. In other words, even some flat RL simulations showed positive slopes, solely due to noise. And many hierarchical RL simulations did not show positive slopes because their underlying parameters prohibited learning.

Bayes Factors take these factors into account when comparing models quantitatively. In this section, we aim to understand how each model produced different markers of hierarchical behavior, expecting systematic links between

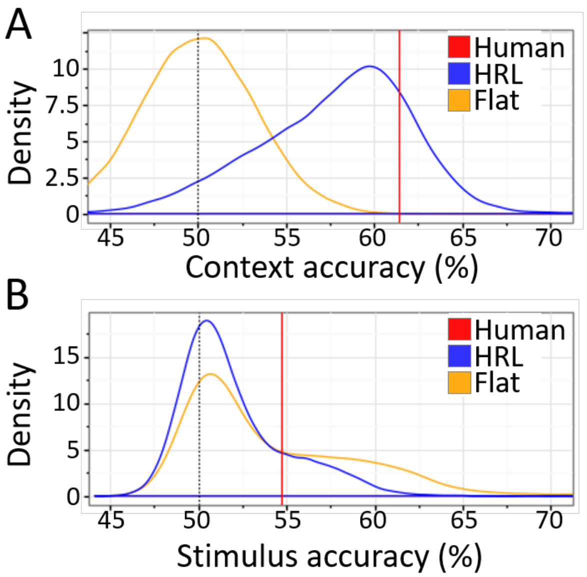

Figure 7: A)-B) Distribution over model behavior in the comparison phase. A) Accuracy for the context condition, i.e., % of trials in which the higher-valued context was chosen. B) Same for stimulus condition.

certain parameters and model behaviors for model that provides a mechanism to explain the result (e.g., positive slopes in the hidden-context phase for hierarchical RL), and the lack thereof when the model cannot.

In the flat model, the forget parameter $f$ influenced the frequency of task-set perseveration errors (suppl. Fig. 9, "Task-set perseveration errors"), as well as overall accuracy (suppl. Fig. 9, "Accuracy trial 1"), as expected. Nevertheless, the levels of task-set perseveration errors never went above, and accuracy never dropped below chance, such that these behaviors did not provide evidence for systematicity, and hierarchy. Extreme values of $f$ also influenced performance differences between context and stimulus condition in the comparison phase (suppl. Fig. 9, "Context minus stimulus"), which could be interpreted as a sign of hierarchy, whereby larger learning rates were related to bigger spreads in this measure.

In the hierarchical model, the forget parameter $f_a$ played a similar role for "Accuracy trial 1" and "Context minus stimulus", but its role for "Task-set perseveration errors" was reversed. $f_a$ also influenced "Task-set reactivation" (suppl. Fig. 8. Other model parameters showed additional relationships with behavioral markers in this model, such that, for example, very small high-level learning rates $\alpha_{TS}$ were associated with more task-set perseveration errors and smaller effects of task-set values on performance ("Effect of task-set values"), and high-level beta $\beta_{TS}$ was associated with increased task-set reactivation, reduced task-set perseveration errors, and increased effect of task-set values on performance. This shows that in the hierarchical model, behavioral markers of hierarchical behavior arose from a complex interplay between model parameters.

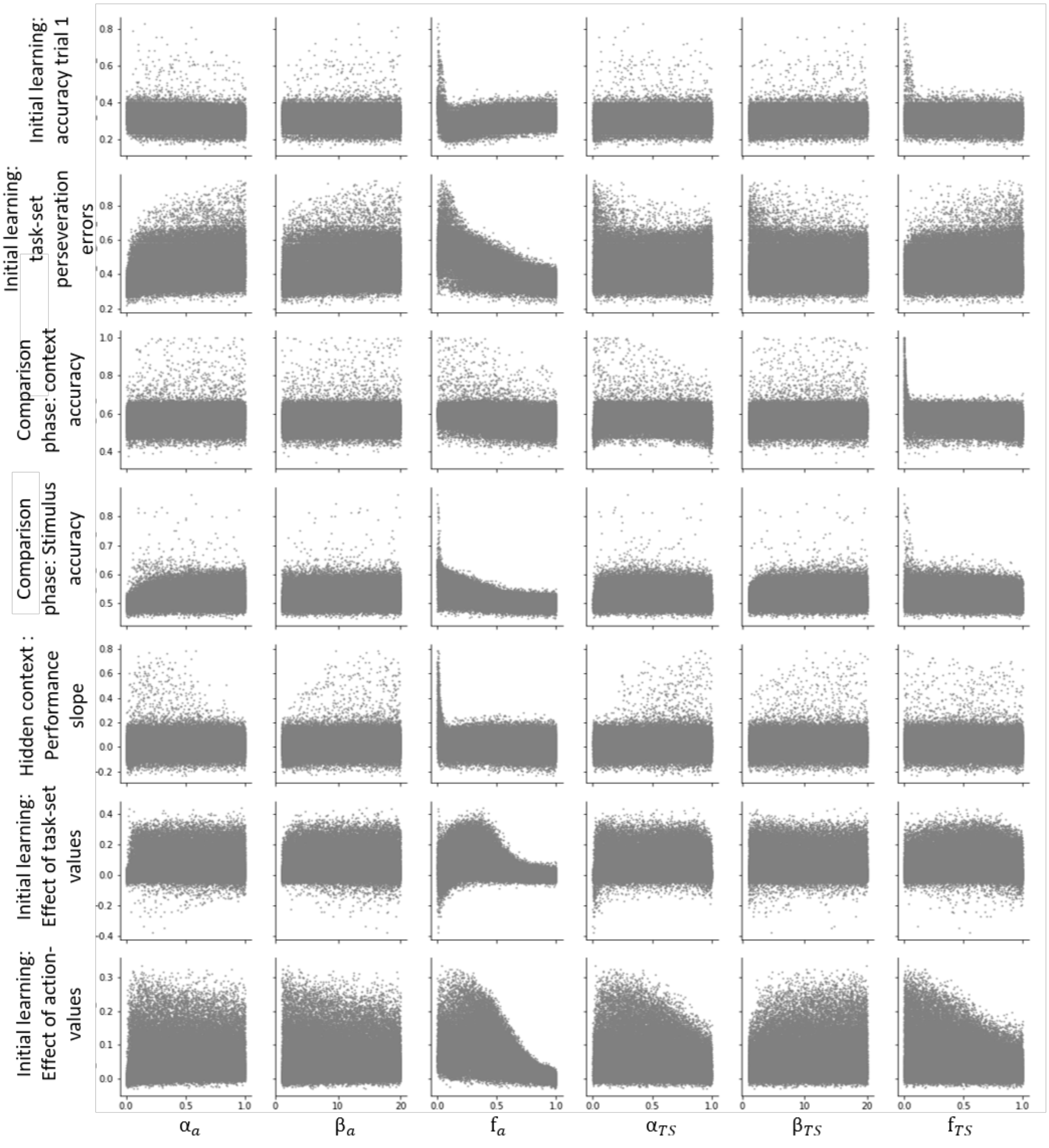

Figure 8: Relationship between hierarchical model parameters and behavioral markers across 50,000 simulations.

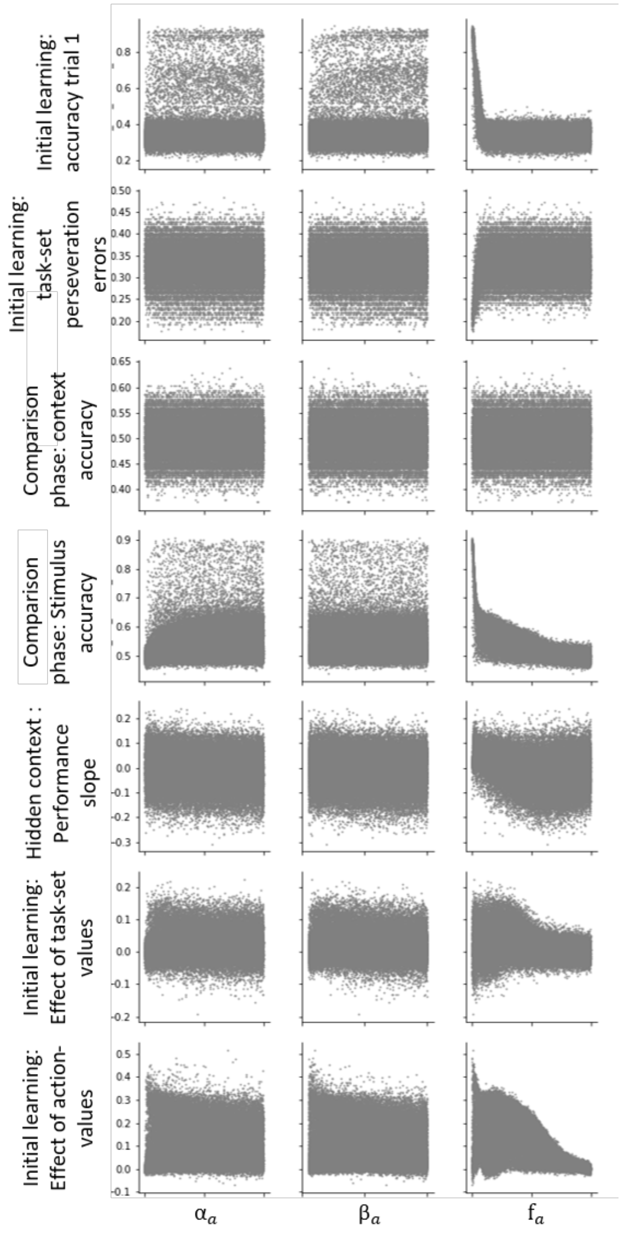

Figure 9: Relationship between flat model parameters and behavioral markers across 50,000 simulations.

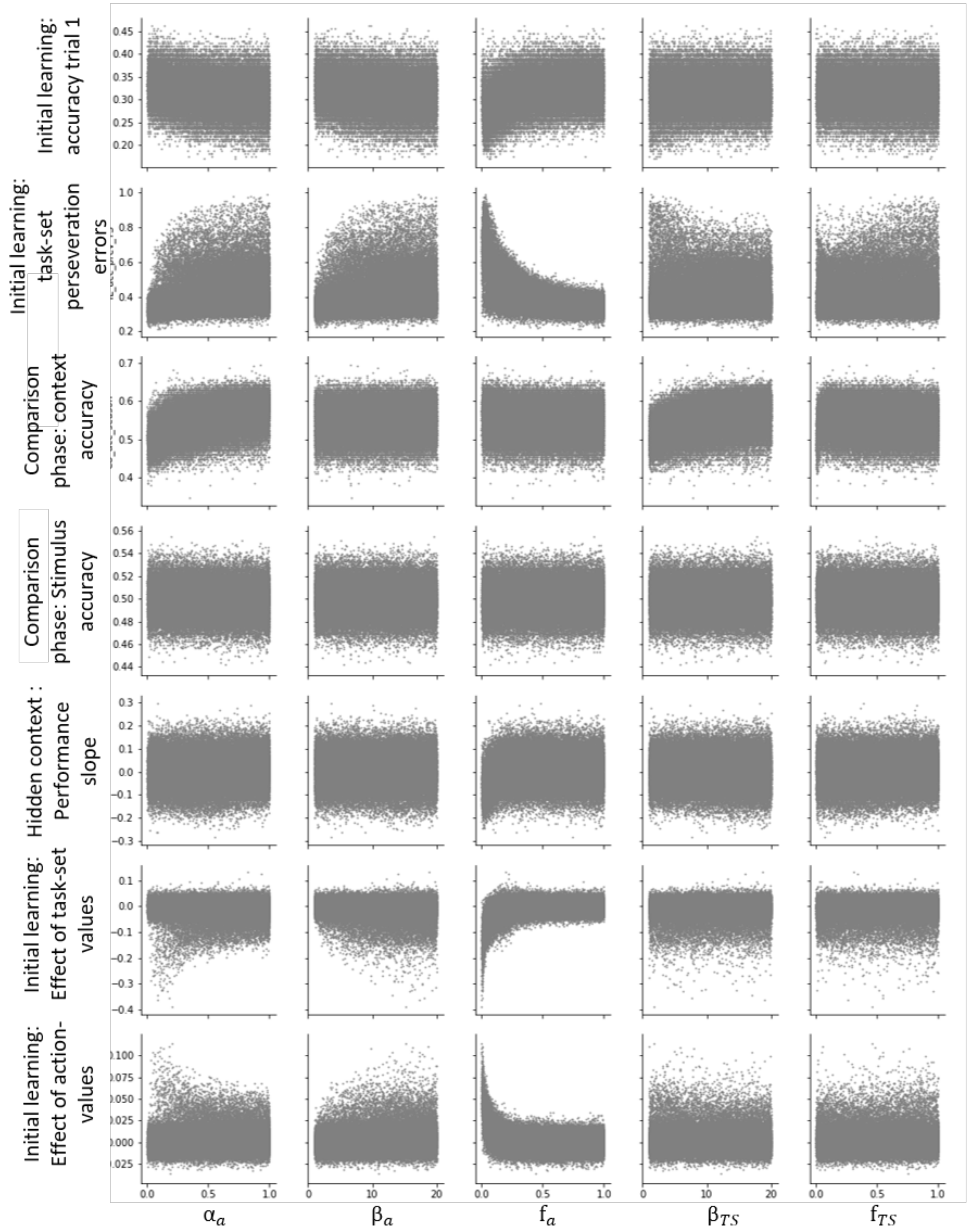

Figure 10: Relationship between hierarchical model parameters and behavioral markers across 60,000 simulations.
